# A novel Pan-viral prophylaxis strategy using vaccine adjuvant CAF09b protects against influenza virus infection

**DOI:** 10.1101/2021.10.21.465243

**Authors:** Julie Zimmermann, Signe Tandrup Schmidt, Ramona Trebbien, Rebecca Jane Cox, Fan Zhou, Frank Follmann, Gabriel Kristian Pedersen, Dennis Christensen

## Abstract

The SARS-CoV-2 pandemic caused a massive health and societal crisis, although the fast development of effective vaccines reduced some of the impact. To prepare for future pandemics, a pan-viral prophylaxis could be used to control the initial virus outbreak in the period prior to vaccine approval. The liposomal vaccine adjuvant CAF^®^09b contains the TLR3 agonist polyinosinic:polycytidylic acid, which induces a type I interferon (IFN-I) response and an antiviral state in the affected tissues. When testing CAF09b as a potential pan-viral prophylaxis, we observed that intranasal administration of CAF09b to mice resulted in an influx of innate immune cells into the nose and lungs and upregulation of IFN-I related gene expression. When CAF09b was administered prior to challenge with mouse-adapted influenza A/Puerto Rico/8/1934 virus, it protected from severe disease, although virus was still detectable in the lungs. However, when CAF09b was administered after influenza challenge, the mice had a similar disease course to controls. In conclusion, CAF09b may be a suitable candidate as a pan-viral prophylactic treatment for epidemic viruses, but must be administered prior to virus exposure to be effective.

## INTRODUCTION

Since the beginning of the 21^st^ century, the world has battled a series of major crises caused by viral epidemics, including SARS-CoV-1, influenza A H5N1 and pandemic H1N1, MERS-CoV, Ebola virus, and the current SARS-CoV-2, which has underscored how vulnerable we are to emerging viral threats. In a breakthrough for mRNA-based vaccines, several vaccines were approved for human use in less than a year after a global pandemic was declared, which is unprecedented and beyond the most optimistic initial hopes (1). These vaccines are very effective at preventing disease and death caused by the current SARS-Cov-2 strains, and are key components to control the pandemic. However, the damage caused by SARS-Cov-2 in the period prior to vaccine approval in terms of human and economic losses emphasizes the need for a ready-to-use tool for immediate control of viral outbreaks.

Most pathogens capable of causing a pandemic have a high mutation rate, which results in evolution into different lineages characterized by variable levels of virulence and transmissibility. This can also lead to emergence of new variants and potentially evasion from vaccine induced immunity as well as disease-acquired protection (2). Several mutations in the immunodominant SARS-CoV-2 Spike protein have been identified, which cause continued concern for the efficacy of the currently licensed vaccines (2, 3).

There is therefore an unmet global need for a strategy that can be implemented immediately after an outbreak to reduce risk of infection, improve disease outcomes and inhibit transmission to other people, without the use of large-scale quarantines, travel restrictions, and social distancing. An optimal strategy would be pan–viral and its efficacy should not be affected by virus homoplasticity. A pan-viral prophylactic strategy against pandemic threats requires protection against a broad and largely unforeseen spectrum of viruses. The innate immune system forms the bodies emergency preparedness against pathogenic threats and has evolved to recognize and react instantly to pathogenic fingerprints like viral double stranded (ds)RNA (4). For the last 20 years, research has identified the mechanisms behind how the innate immune system reacts to gain initial control of infections, until the adaptive immune system has had time to form a specific immune response to eliminate the invading pathogen. The innate immune system is therefore in most cases not sufficient to eliminate viral infections with e.g. influenza viruses or coronaviruses, but it can mitigate damaging effects of the virus attack and reduce disease symptoms if properly activated.

One of the key components of the early innate response against viruses is type I interferons (IFN-I). The timing of initiation of an IFN-I response depends on viral and host factors and is critical for the progression of a SARS-CoV-2 infection (5). Thus, an initial low viral load allows the rapid induction of a strong IFN-I response, which can clear the infection, whereas an initial high viral load will suppress the IFN-I response and cause disease progression (5). Supporting this, *in vitro* models show that pre-treatment with IFN-α and IFN-β effectively prevents infection with SARS-CoV-2 upon challenge (6). Furthermore, the IFN-I responses were impaired in SARS-CoV-2 infected patients with severe and critical disease (7). IFN-I are pleiotropic immunomodulatory cytokines that can activate protective antiviral effects in several cell types, which in concert induce a general antiviral state (8). Danger signaling is initiated when viral RNA is detected by pattern recognition receptors (PRRs) e.g. the RIG-I, TLR3, −7 and −8 receptors activating IFN-I secretion. However, due to the effective antiviral effects induced by IFN-I, viruses have developed several ways to circumvent an antiviral state, mainly through blocking expression of IFN-I related genes (9).

Modern pharmaceutical design and engineering has made it possible to synthetically design TLR agonists and, combined with novel delivery techniques, these can now be delivered safely to mucosal surfaces, which permits clinical testing. In recent clinical trials intranasal delivery of the TLR3 agonist polyinosinic:polycytidylic acid [poly(I:C)], a synthetic dsRNA structurally similar to virus dsRNA, stimulated IFN-I production and significantly protected against Rhinovirus and Influenza virus infections in humans (10). This immune prophylaxis was safe and well tolerated. Infection was less severe, had shorter duration and importantly reduced the number of individuals with study-defined, laboratory confirmed illness, without compromising seroconversion induced by infection. The novel vaccine adjuvant CAF^®^09b combines the TLR-3 agonist poly(I:C) with C-type lectin receptor agonist monomycoloyl glycerol (MMG), formulated in a cationic liposomal delivery system and was developed for vaccines against viral infections (11, 12) and cancer (13). CAF09b has thus been combined with numerous vaccine antigens and tested extensively as vaccine adjuvant preclinically and in humans (NCT03412786, NCT03715985) (14). CAF09b solves a range of challenges related to the toxicity associated with administration of therapeutic doses of poly(I:C) (15, 16). When poly(I:C) is complexed within cationic liposomes, the detrimental innate immune reactions are abrogated (16).

In the present study, we show that intranasal administration of CAF09b to mice caused upregulation of several IFN-I related genes, and pretreatment with CAF09b prevented death upon lethal challenge with mouse-adapted influenza A A/Puerto Rico/8/1934 (H1N1) (PR8) virus. In contrast, no protective effect was observed when CAF09b treatment was initiated after influenza challenge. Thus, CAF09b is a promising tool for pan-viral prophylaxis against viruses with pandemic potential.

## METHODS

### Preparation of CAF09b^®^

Dimethyldioctadecylammonium bromide (DDA) and monomycoloyl glycerol (17) were obtained from NCK A/S (Farum, Denmark) and polyinosinic:polycytidylic acid was bought from Dalton Pharma Services (North York, Canada). The liposomal adjuvant CAF09b was prepared essentially as described elsewhere (13). Briefly, weighed amounts of DDA and MMG were dissolved in EtOH, 96%. A lipid film was formed by evaporating the EtOH under a gentle N2 stream for 2h followed by air-drying overnight. The lipid film was rehydrated in Tris-buffer (10 mM, pH 7.0) with 2% w/v glycerol by high shear mixing by using a Heidolph Silent Crusher equipped with a 6F shearing tool (Heidolph Instruments GmbH, Schwabach, Germany) at 60 °C and 26,000 rpm for 15 min. Poly(I:C) was added continuously during high shear mixing using a peristaltic pump (Pharmacia Biotech, Stockholm, Sweden) over 30 min. The final CAF09b dose was 250/50/12.5 μg DDA/MMG/poly(I:C).

### In vivo studies

The induction of innate immune cell responses and prevention of influenza disease after intranasal administration of CAF09b was evaluated *in vivo* in CB6F1 mice (BALB/c x C57BL/6, Envigo, Horst, The Netherlands). The animal experiments were conducted in accordance with EU directive 2010/63/EU and regulations set forth by the Danish National Committee for the Protection of Animals used for Scientific Purposes. The mice were randomized in the study groups (n=6 or 8) and allowed free access to food, water and recreational stimuli.

The mice were treated with 20 μl CAF09b i.n. or Tris-buffer (10 mM, pH 7.0) as a negative control at different time points prior to termination (the day after the last treatment) or to challenge with 30 μl influenza A Mouse-adapted influenza virus strain A/Puerto Rico/8/1934 virus (5×10^3^ EID_50_/ml administered as 15 μl/nostril). The virus was propagated in the allantoic cavity of 10-day-old embryonate hen’s eggs. Allantoic fluid was harvested, clarified and frozen at −80 degree C until used. In the influenza virus challenge studies, the mice were followed for 7 days after challenge and monitored for change in weight and disease score (ruffled fur, neurological signs and respiratory symptoms). The mice were euthanized during the course of the study if they met the predefined humane endpoints; weight loss exceeding 20% of initial weight, a disease severity score of 2 for more than 48 h or 3 for more than 12 h. At termination one lung was removed into RNAlater (Thermo Fisher Scientific, Waltham, MA, USA) for evaluation of induction of IFN-I, and the other lung was removed into RPMI 1640 (Gibco, Invitrogen, Carlsbad, CA, USA) for assaying the infectious influenza virus titer. Blood was collected for evaluation of PR8 H1N1-specific antibody titers. In the study terminated prior to influenza virus challenge, the mice were administered anti-CD45:FITC intraveneously 3 min. prior to euthanization for staining of blood leukocytes. After euthanization, the nasal tissue (upper jaw and nose in front of the eyes) and lungs were removed into RPMI 1640 for identification of the innate cells responses.

### Innate immune cell characterization by flow cytometry

For evaluation of the innate cell response in the lungs and noses after treatment with CAF09b, the lungs and noses were processed to obtain single cell suspensions. Each lung was immersed in 2.5 ml cRPMI (RPMI 1640 supplemented with 5 × 10 M 2-mercaptoethanol, 1% pyruvate, 1% HEPES, 1% (v/v) premixed penicillin-streptomycin solution (Invitrogen Life Technologies), 1 mM glutamine, and 10% (v/v) fetal calve serum (FCS)) with 1.6 mg collagenase (Sigma-Aldrich, St. Louis, MO, USA) and processed twice on a GentleMacs using the lung program (Miltenyi Biotec, Köln, Germany) with 30 min incubation at 37°C in between. The homogenate was then passed through a 100 μm nylon mesh cell strainer (Corning Inc., Corning, NY, USA) and washed twice in cold PBS (Gibco, Invitrogen, Carlsbad, CA, USA). The noses were cut into smaller pieces prior to incubation with 1.6 mg collagenase in 2.5 ml cRPMI+10% FCS for 30 min at 37°C with agitation. The detached cells were then passed through a cell strainer and washed twice in cold PBS. The single cell suspensions were placed in 96-well V-bottomed plates at 10^6^ cells/well, treated with Fc-block and stained with live-dead cell marker:AF488, CD19:FITC, Ly6G:PE, CD49d:PerCP-Cy5.5, CD11b:PE-Cy7, F4/80:APC, Ly6C:APC-Cy7, NK1.1:BV421, CD11c:BV510 and MHC II (IA-IE):BV605 (eBiosciences, San Diego, CA, USA or BD Biosciences, San Jose, CA, USA). The cells were analysed using a LSRFortessa with FACSDiva software (BD Biosciences) and the data was analysed using FlowJo (BD Biosciences). Cells were identified as macrophages (F4/80^+^,CD11b^+^), neutrophils (Ly6G^+^), natural killer (NK) cells (NK1.1^+^), monocytes (CD11b^+^,Ly6C^+^) and dendritic cells (DC) (MHCII^+^,CD11c^+^), the gating strategy is shown in Supplementary Fig. 1.

### Type I IFN induction by qPCR

Lungs were removed into RNAlater where they were kept at 4°C for minimum 24h and then stored at −20° C. RNA was isolated using RNeasy mini kit (Qiagen, Hilden, Germany), lung tissue was homogenized by gentleMACS using the RNA_01 program (Miltenyi Biotec). Genomic DNA was removed by on-column DNase digestion using RNase-Free DNase set (Qiagen, Hilden, Germany). The quality of the RNA was determined by NanoDrop™ 2000/2000c Spectrophotometers (Thermo Fisher Scientific, Waltham, MA, USA) and 2100 Bioanalyzer (Agilent Technologies, Santa Clara, CA, USA). All samples had a RIN value > 8. The cDNA was synthesized by RT^2^ First Strand Kit (QIAGEN, Hilden, Germany). Quantitative real-time PCR (qRT-PCR) was performed on LightCycler® 480 (Roche, Basel, Schweiz) using AbsQuant 2nd Derivative Max for obtaining the Ct value. PCR conditions were 10 min 95°C followed by 45 2-step cycles of 15 s at 95°C and 1 min 60°C. For RNA profiling, the RT^2^ Profiler Array “Type I Interferon Response” (Cat. No. 330231 PAMM-016ZA) (Qiagen, Hilden, Germany) was used. The relative mRNA amount was obtained by the ΔΔCt method (18), with the use of the three housekeeping genes GAPDH, GUSB and HSP90AB1.

### Virus titer determination by qPCR

The virus titers were determined on lung supernatants from infected mice. The lungs were removed into RPMI and homogenized by gentleMACS using the RNA_01 program (Miltenyi Biotec, Köln, Germany). The lung supernatant was stored at −80°C, and the RNA was isolated by Quick-RNA Viral Kit (Zymo Research, Tustin, CA, USA). The quality of the RNA was determined using a NanoDrop™ 2000/2000c Spectrophotometer (Thermo Fisher Scientific). The titers were determined by the virotype Influenza A RT-PCR Kit (Indical Bioscience, Leipzig, Germany), using 70ng of RNA. The qRT-PCR was performed on LightCycler® 480 (Roche, Basel, Schweiz) using AbsQuant 2nd Derivative Max for obtaining the Ct value. PCR conditions were 10 min at 45°C, 10 min at 95°C following 40 cycles of 15 s at 95°C and 1 min at 60°C. The relative mRNA amount was obtained by the ΔΔCt method (18), using β-actin as housekeeping gene.

### PR8 H1N1 hemagglutinin protein-specific antibody ELISA

Serum was obtained after centrifugation of the blood for 10 min at 10,000*g* and stored at - 20°C until further use. The PR8 H1N1 antigen-specific IgG antibody responses induced by influenza PR8 challenge were evaluated by ELISA. PR8 H1N1 hemagglutinin protein (Sino Biological Inc., Beijing, China) 1 μg/ml was coated onto MaxiSorp plates (Nunc, Hillerød, Denmark) overnight at 4°C. Serum was added at 10-fold serial dilutions and incubated for 2 h at room temperature (rt) followed by incubated with HRP-conjugated anti-mouse total IgG antibodies (AH diagnostics, Tilst, Denmark) for 1 h at rt. The signal was detected by TMB (Kem-En-Tec, Taastrup, Denmark) and the reaction stopped with 0.2M H_2_SO_4_ followed by analysis on a TECAN Sunrise™ ELISA reader (Tecan Trading AG, Männedorf, Schweiz) at 450 nm with 620 nm correction.

### Statistical analysis

Statistical analyses were performed using either GraphPad Prism software version 8.3.0 for Windows (GraphPad Software, La Jolla, CA, USA) or R (version 4.0.2). Statistical significance between multiple groups was determined by one-way ANOVA followed by either Dunnett’s multiple comparisons test (if all groups are compared to the naïve) or Tukey’s multiple comparisons test (if all groups are compared to each other). Statistical significance between two groups was determined by two-tailed unpaired t-test. Two fold up/down regulated genes were determined in R and plotted in a scatterplot (car package). The difference in relative mRNA expression for each genes was calculated by the z-score and illustrated in a heatmap (pheatmap package).

## RESULTS

### Intranasal delivery of CAF09b upregulates IFN-I related genes

The ability of CAF09b to induce IFN-I related genes was evaluated in a mouse model. CAF09b was administered i.n. twice on days 0 and 3 as well as daily (days 0, 1, 2, and 3). A naïve group was administered Tris-buffer on days 0, 1 and 3. The induction of IFN-I related genes in the lungs was analysed by qPCR on day 4. Intranasal administration of CAF09b upregulated several IFN-I related genes (Fig. 1A, B). Out of the 84 IFN-I related genes assayed, 33 and 42 genes were more than 2-fold upregulated compared to naïve mice after administration of two and four doses of CAF09b, respectively. No genes were downregulated more than 2-fold after administration of CAF09b compared to naïve mice (Fig. 1A, Supplementary Fig. 2).

**Figure 1:**
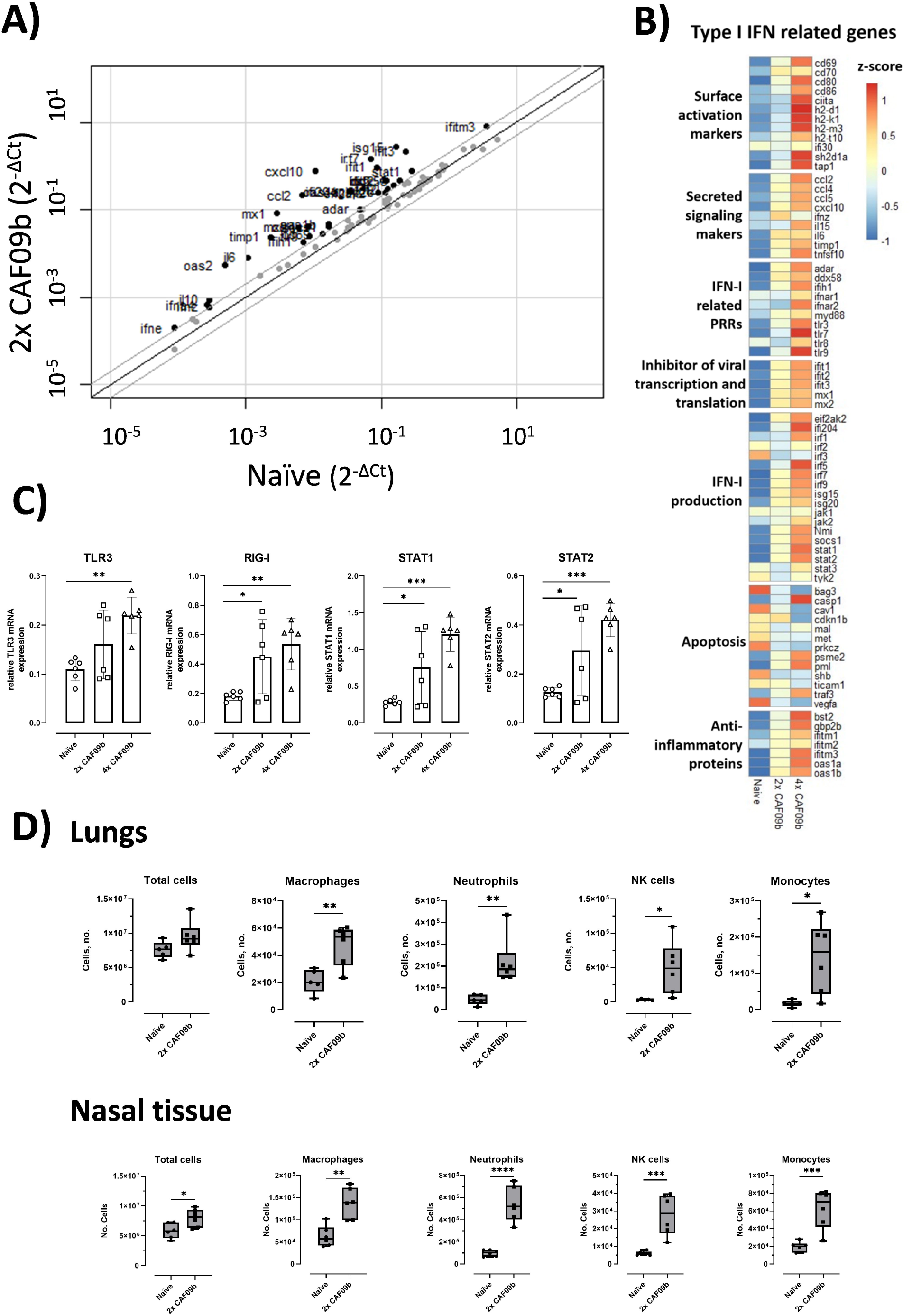
Intranasal administration of CAF09b upregulates IFN-I related genes and causes an influx of innate immune cells. Mice (CB6F1, n=6) were administered CAF09b 20 μl i.n. on days 0, 3 (2x CAF09b) or days 0, 1, 2, 3 (4x CAF09b), the naïve group was administered Tris-buffer 20 μl i.n. on days 0, 1, 3. (A-C) Lungs were collected at day 4 and IFN-I responses were measured by qPCR. **(A)** Scatterplot of the 84 genes related to IFN-I responses. Naïve mice where compared to 2x CAF09b treated mice. The black line represents 1 fold change and grey lines represent 2 fold change. Dots represent the average relative mRNA expression for each of the genes. **(B)** Heatmap of an average of the relative mRNA expression for all 84 genes. The three groups are compared against each other and represented by the z-score. **(C)** Plots of the relative mRNA expression value for the four genes TLR3, RIG-I, STAT1, STAT2. Each dot represent one mouse, boxes denote mean±S.D. One-way ANOVA followed by Dunnett’s multiple comparisons test with a comparison to the naïve group, * p-value ≤ 0.05, ** p-value ≤ 0.01, *** p-value ≤ 0.001. **(D)** The cellular composition in the lungs and nasal tissue was analyzed at day 4. The total amount of cells in each tissue was counted, while macrophages (F4/80^+^,CD11b^+^), neutrophils (Ly6G^+^), NK cells (NK1.1^+^), monocytes (CD11b^+^,Ly6C^+^) and DCs (MHCII^+^,CD11c^+^) were identified by flow cytometry (Supplementary Fig. 1). Box and whiskers plots denoting mean and min./max. value, dots represent individual mice. Two-tailed unpaired t-test, * p-value ≤ 0.05, ** p-value ≤ 0.01, *** p-value ≤ 0.001.

Poly(I:C) is a ligand for TLR3 in endosomes (19), and it is generally believed that RIG-I is one of the primary receptors of cytoplasmic dsRNA (20). TLR3 was significantly upregulated in the group administered four doses of CAF09b compared to in the naïve group, while RIG-I was significantly upregulated after administration of both two and four doses of CAF09b compared to the naïve group (Fig. 1C). STAT1 and STAT2 are transcription factors of Interferon Stimulated Genes (ISGs) and key elements in the IFN-I response. Both two and four doses of CAF09b significantly upregulated STAT1 and STAT2 transcription (Fig. 1C).

### Intranasal administration of CAF09b induces influx of several innate immune cell subsets

Prophylactic local administration of CAF09b was shown to prevent death after challenge with mouse-adapted PR8 influenza. In addition to the induction of IFN-I observed in the lungs, we evaluated the influx of different innate immune cell subsets into the lungs and nasal tissue, respectively, following i.n. administration of two doses of CAF09b (Fig. 1D). The total cell count in the lungs was not significantly increased by administration of CAF09b, while a significant increase in total cells was observed in the nasal tissue. However, an increase in innate immune cell subsets was observed in both organs compared to naïve mice, although of different magnitude. Thus, the levels of macrophage (F4/80^+^,CD11b^+^), neutrophil (Ly6G^+^), NK cell (NK1.1^+^), monocyte (CD11b^+^,Ly6C^+^) and DC (MHCII^+^,CD11c^+^) subsets were all increased in the lungs and nasal tissue of mice after two doses of CAF09b.

### Two doses of CAF09b protected against influenza disease

The ability of CAF09b to protect against influenza challenge was evaluated in the murine model of influenza A H1N1 PR8 virus infection. Two different dosing regimens were tested and thus CAF09b was administered i.n. twice (on days −6 and −3 prior to challenge) or four times (days −6, −5, −4, and −3). In the mock group, the mice were administered Tris-buffer twice on days −6 and −3. Mice were challenged with mouse-adapted PR8 influenza (150 EID_50_, 30 μl i.n.) on day 0 and their survival monitored for 7 days post influenza challenge (p.i.c.) (Fig. 2). All mice administered two doses of CAF09b survived in the study, whereas 6/8 and 3/8 mice survived in the groups administered four doses of CAF09b or Tris-buffer, respectively. Thus, two doses of intranasal CAF09b protected against severe influenza disease and there was no additional benefit on disease outcome from using additional CAF09b doses. It was therefore decided to continue the studies with two CAF09b administrations.

**Figure 2:**
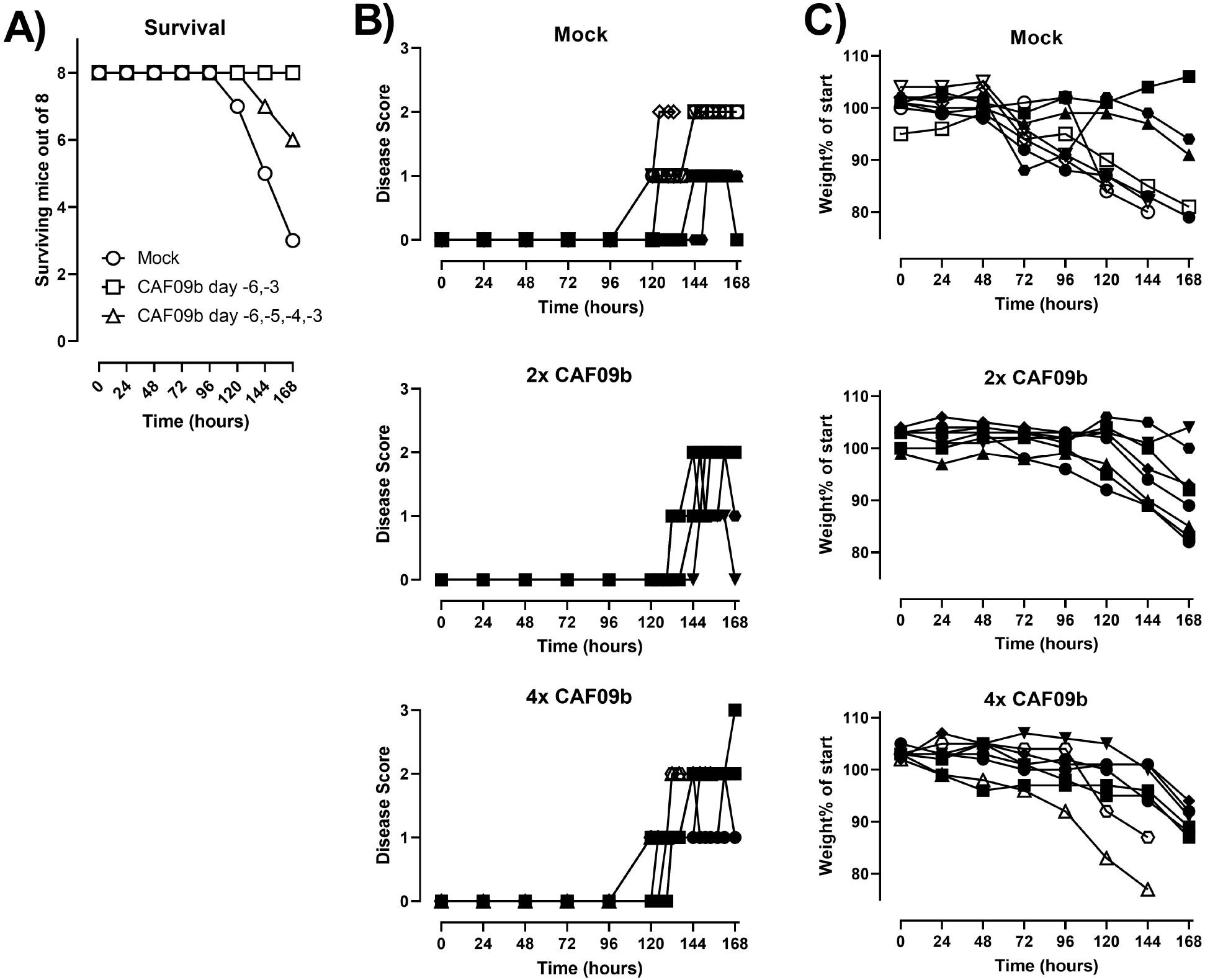
CAF09b i.n. protects against influenza disease. Mice (CB6F1, n=8) were administered CAF09b 20 μl i.n. on days −6, −3 (2x CAF09b) or days −6, −5, −4, −3 (4x CAF09b), the mock group was administered Tris-buffer 20 μl i.n. on days - 6, −3. The mice were challenged with mouse-adapted influenza A PR8 H1N1 virus (150 EID_50_, 30 μl i.n.) on day 0. **(A)** Survival curves, mice were euthanized when meeting humane endpoints. **(B)** The disease score was monitored for 168 h after influenza challenge. **(C)** The body weight as percentage of initial weight (measured on day −1) was measured for 168 h. (B,C): Open symbols; mice were euthanized prior to study termination, closed symbols: mice were euthanized at study termination.

### CAF09b is only effective when administered before influenza challenge

In a pandemic scenario using CAF09b as viral prophylaxis, the exact period between administration of the adjuvant and encountering the pathogen would be unknown. Therefore, we evaluated the effect of CAF09b on preventing disease and death after influenza challenge at different periods around CAF09b administration. Thus, two doses of CAF09b were administered i.n. at days −11 and −8 before influenza challenge (b.i.c.), −5 and −2 b.i.c., −2 b.i.c. and +1 p.i.c., and +1 and +4 p.i.c. A mock group was administered Tris-buffer twice on day −5 and −2 (Fig. 3A). The disease severity score, body weight and survival were evaluated over 7 days p.i.c. (Fig. 3B, C). For the groups treated with CAF09b prior to influenza challenge, disease symptoms were reduced but not completely prevented and the onset of disease, measured as an increase in disease severity score, occurred later than for the mock group (Fig. 3B, C). In contrast, the disease symptoms and weight curves of the mice starting treatment after influenza challenge were similar to the mock group.

**Figure 3:**
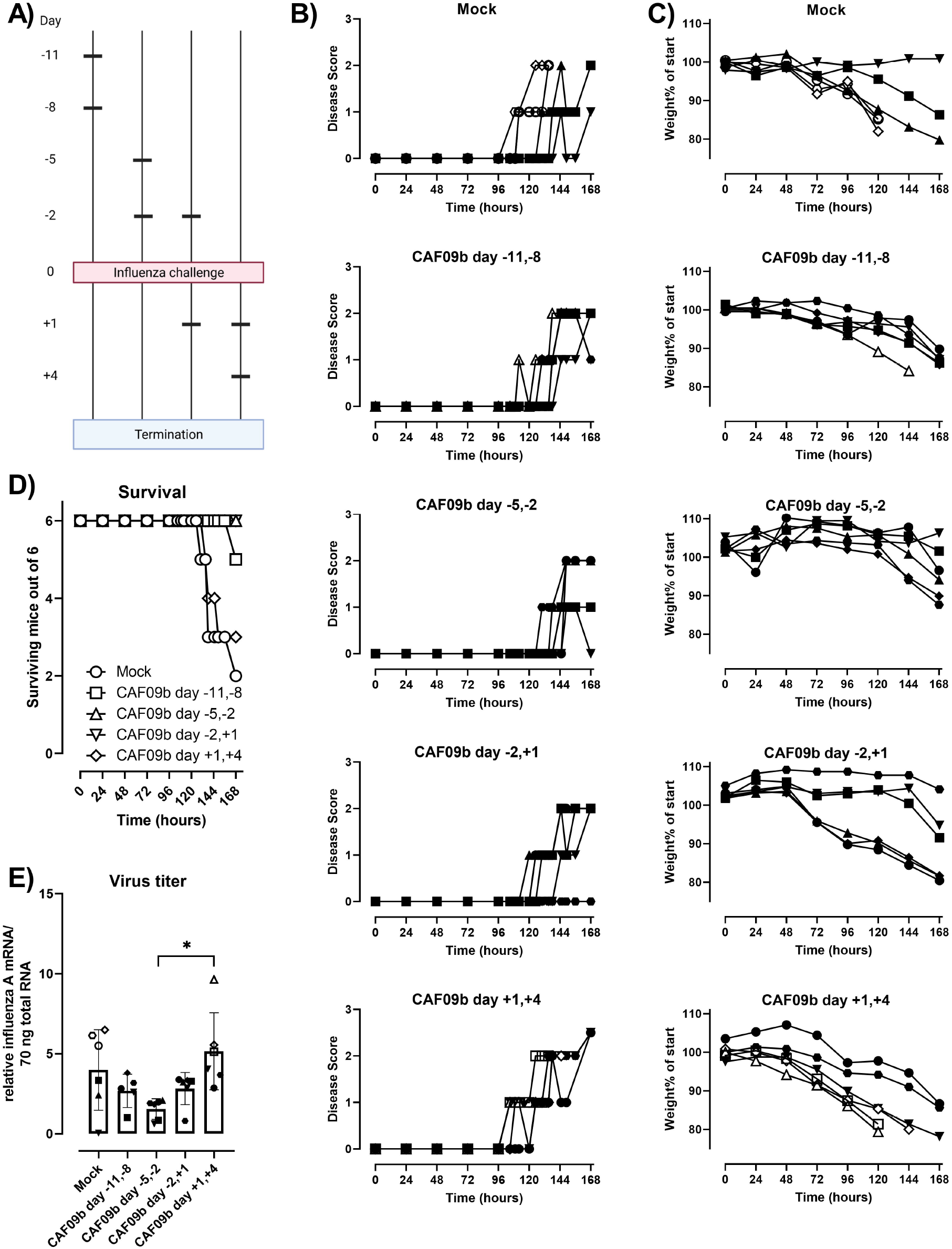
The number of CAF09b administrations affects protection against influenza virus challenge. **(A)** Mice (CB6F1, n=6) were administered CAF09b 20 μl i.n. twice on days −11,-8; −5,-2; - 2,+1 and +1,+4 and PR8 influenza virus challenge (150 EID_50_, 30 μl i.n.) on day 0. Tris-buffer 20 μl administered i.n. on day −5 and −2 was used as a negative control (mock). **(B)** The disease score was monitored for 168 h after influenza challenge. **(C)** The body weight as percentage of initial weight (measured on day −1) was measured for 168 h. **(D)** Survival curves, mice were euthanized when meeting humane endpoints. **(E)** The PR8 virus titer was determined in the lungs at the point of euthanization by qPCR, boxes denote mean±S.D. One-way ANOVA followed by Tukey’s multiple comparisons test, * p-value ≤ 0.05, ** p-value ≤ 0.01 (B,C,E): Open symbols; mice were euthanized prior to study termination, closed symbols: mice were euthanized at study termination. (A) was created with Biorender.

The survival after influenza challenge correlated with the disease severity scores and rate of weight loss. Thus, 6/6 or 5/6 mice survived the study when CAF09b treatment was initiated prior to influenza challenge (Fig. 3D). In contrast, 2/6 mice survived in the mock group and 3/6 mice survived when CAF09b treatment started after influenza challenge. CAF09b treatment should therefore be initiated prior to influenza virus infection to alleviate disease and improve survival. The virus titers in mice at the termination of the study were similar across the groups irrespective of CAF09b treatment, although there was a tendency towards lower influenza titers in the day −5, −2 b.i.c group (Fig. 3E)

The expression of genes related to IFN-I responses was measured 7 days p.i.c. in mice administered CAF09b at days −5 and −2 b.i.c; −2 b.i.c. and +1 p.i.c., and +1 and +4 p.i.c. as well as the mock group. The influenza infection highly influenced the expression of IFN-I related genes. Out of the 84 IFN-I related genes, 10-14 genes were more than 2-fold up- or down regulated when compared to the mock group. When comparing the naïve group to the mock group, 53 genes where more than 2-fold up or down regulated and 42 of these were only different in the naïve vs mock (Fig 4A, Supplementary Fig 3). When focusing on the 14 highly up- or down regulated genes there was a similar expression profile among the groups treated with CAF09b. CAV1, CRP, MET, PRKCZ and VEGFA gene expression was lower in the CAF09b treated groups compared to the mock group, although not statistically significant. In contrast, CCL5 was significantly upregulated in the CAF09b day −5, −2 group compared to both mock and the group that had received CAF09b at day −2 b.i.c. and +1 p.i.c., where in contrast both CXCL10 and IL6 were significantly upregulated compared to in the mock group (Fig 4B).

**Figure 4:**
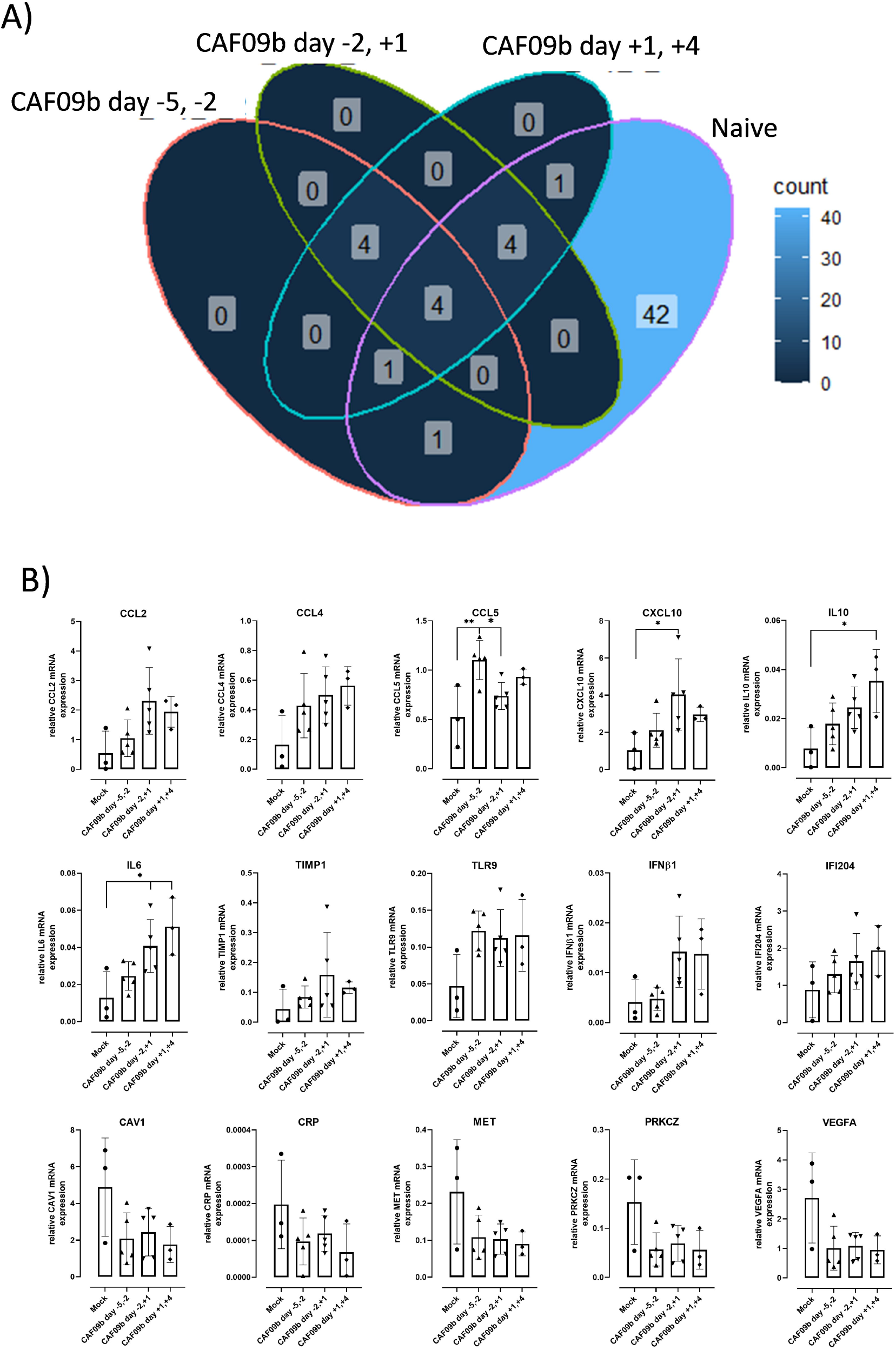
Timing of CAF09b administration minimally affects infection induced gene expression. Mice (CB6F1, n=3-6) were administered 20 μl CAF09b i.n. twice on days −5,-2; −2,+1 and +1,+4 and challenged with PR8 influenza virus (150 EID_50_, 30 μl i.n.) on day 0. A naïve and a mock group were administered 20 μl Tris-buffer i.n. on day −5 and −2, and the mock group was challenged with PR8 influenza virus (150 EID_50_, 30 μl i.n.) on day 0. Lungs were collected 7 days post virus challenge and 84 genes related to IFN-I responses were measured by qPCR (lungs were taken at day −1 in the naïve group). **(A)** The Venn diagram shows the number of genes that are more than 2 fold up or down regulated compared to the mock group for the four groups (CAF09b day −5, −2; CAF09b day −2, +1; CAF09b day +1, +4; naïve (Supplementary Fig. 3). **(B)** Plots of the relative mRNA expression value of the genes where the mean value is more than 2 fold up or down regulated in (A). Each dot represent one mouse, boxes denote mean±S.D. One-way ANOVA followed by Tukey’s multiple comparisons test, * p-value ≤ 0.05.

### Treatment with CAF09b does not prevent induction of an influenza-specific antibody response

Induction of an adaptive pathogen-specific memory immune response after infection is critical for protecting the individual from reinfection. To assess if CAF09b treatment interfered with induction of antibody responses, the levels of PR8 H1N1-specific total IgG antibodies were determined in the blood of the mice at the termination of the study or upon euthanization (Fig. 5). All mice in the study developed PR8 H1N1-specific IgG antibodies, indicating that administration of CAF09b did not prevent the induction of adaptive immune responses, despite reducing viral load and disease.

**Figure 5:**
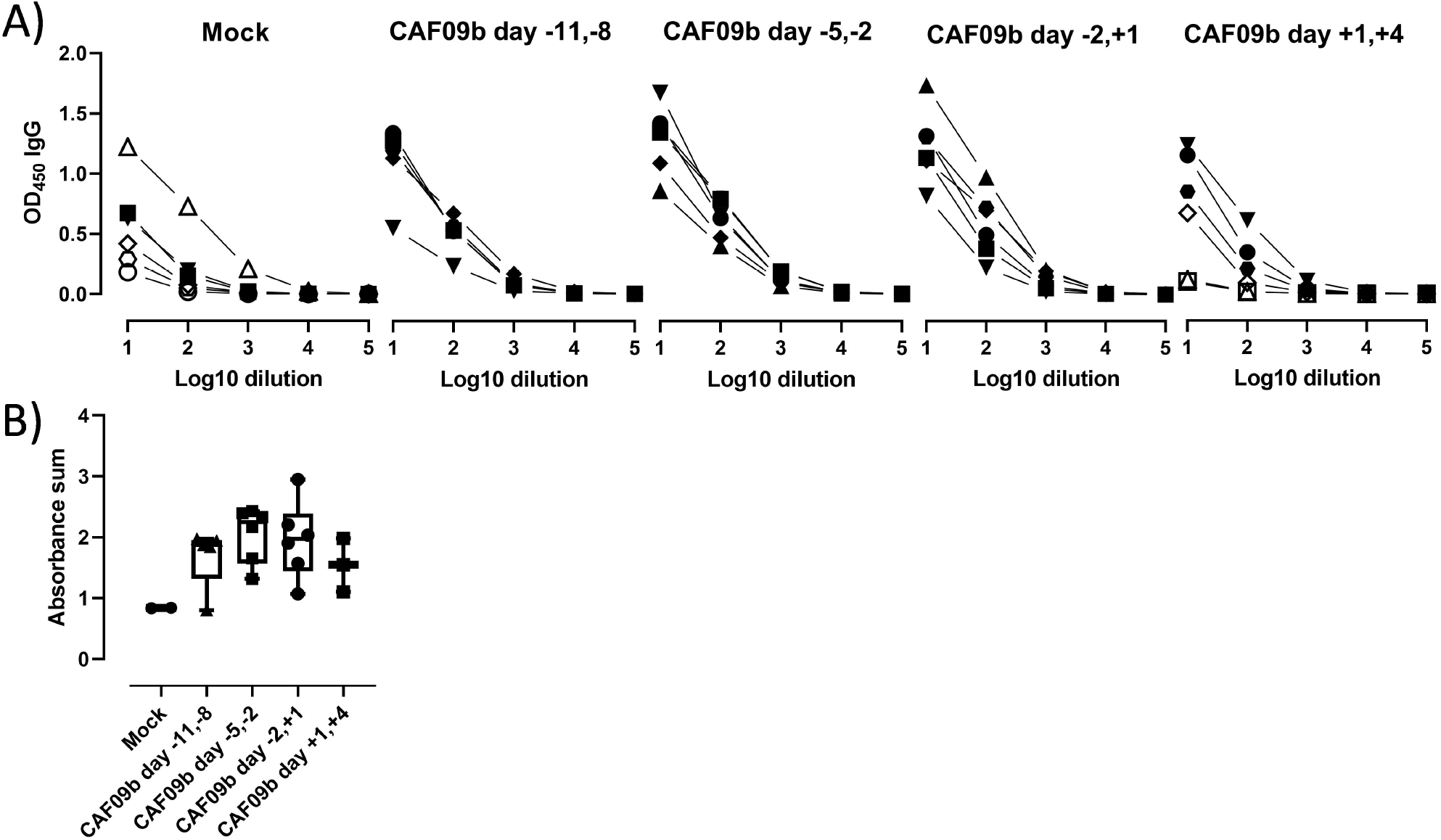
CAF09b does not interfere with induction of antibody responses. Mice (CB6F1, n=6) were administered 20 μl CAF09b i.n. twice on day −11,-8; −5,-2; −2,+1 and +1,+4 and challenged with influenza A PR8 H1N1 virus (150 EID_50_, 30 μl i.n.) on day Tris-buffer 20μl administered i.n. on day −5 and −2 was used as control (mock). **(A)** The PR8-specific total IgG antibodies in serum were determined by ELISA upon euthanize. Open symbols; mice were euthanized prior to study termination, closed symbols: mice were euthanized at study termination. **(B)** The sum of absorbance for the individual mice. Only mice, which were euthanized at study termination are included. Box and whiskers plots denoting mean and min./max. value, dots represent individual mice.

## DISCUSSION

Pan-viral prophylaxis could be an effective first-line measure to prevent a potential viral epidemic or pandemic by providing a readily available, pathogen-nonspecific treatment. Early after identification of a novel virus with epidemic potential, stimulators of innate immunity which effectively induce antiviral responses could be applied to front-line health care workers or close contacts such a household members of infected individuals. We demonstrated here that CAF09b could be used for viral prophylaxis against influenza. The user acceptance of a pan-viral agent however does not only rely on the ability to prevent disease, but also on ease of administration and frequency. A nose spray to be applied 2-3 times a week may well be acceptable until a vaccine is available.

Airway administration of CAF09b recruited innate immune cells and robustly upregulated IFN-I associated genes in the lungs, which together may contribute to the reduction in disease symptoms and prevention of death upon a subsequent influenza infection. The influx of innate immune cells is likely due to the cationic charge of CAF09b, as very similar influx patterns have been observed following intraperitoneal administration of CAF09b and CAF04, a similar adjuvant without poly(I:C) (unpublished data, manuscript in preparation). The cationic nature of the adjuvant causes a local inflammatory response (21), which in turn recruits innate immune cells. The cell populations recruited to the nose and lungs after CAF09b administration may have both beneficial and detrimental effects on the antiviral response. Thus, neutrophils may exert antiviral effects e.g. by secreting antiviral agents such as reactive oxygen species and α-defensins, but may also have damaging effects by promoting a prolonged inflammatory response at the site of infection (22). The role of NK cells in viral infections is not fully understood, but they are recruited in large numbers by different viruses and may contribute to protection both via direct cytotoxicity and via inducing an antiviral state (23).

Early IFN-I responses are critical to prevent disease due to virus infections and prophylactic or therapeutic antiviral strategies aiming to induce IFN-I after intranasal administration have been evaluated in clinical trials. Two PrEP-001 human clinical trials administering powdered poly(I:C) i.n. twice 48 and 24 h prior to challenge with either rhinovirus or influenza A virus showed reduced development of clinical illness and symptoms (10). In another study, IFN-β-1a was intranasally dosed once daily up to 14 days to patients hospitalized with Covid-19 symptoms, which was well tolerated and resulted in greater chances of recovery compared to a placebo group (24). Supporting these findings, a similar clinical study administering IFN-α2b intranasally to patients admitted to hospital with Covid-19 reduced pro-inflammatory cytokine levels and improved the recovery rate compared to treatment with the antiviral agent arbidol hydrochloride (25). However, the timing of direct IFN-I (-α or –β) administration has to be considered, as there is a concern of exacerbation of the cytokine storm observed in later stages of severe disease (24). Indeed, abrogation of IFN-I responses in *Ifnar-/-* Balb/c mice resulted in milder disease after SARS-CoV infection compared to wild type Balb/c mice (26). Furthermore, disease-delayed IFN-I caused inflammation and abrogated the antigen-specific T-cell responses (26).

The importance of an early IFN-I response was demonstrated in studies with the SARS-CoV-1 virus. Here animal studies showed that even in complete absence of T and B cells, animals were able to control the infection if the innate immune system was alert and able to instantly produce IFN-I after infection (27, 28). These studies also showed that the early innate immune responses could facilitate stronger adaptive immunity to infection. In support of this, both prophylactic and post-exposure strategies involving specific innate immune stimulation, especially via TLR3, has been shown to be able to prevent or eliminate a range of viral infections (29, 30).

Importantly, we showed that CAF09b had to be administered prior to influenza challenge to be effective at preventing symptomatic disease and improve survival. This is in accordance with a study using the poly(I:C) analogue Hiltonol^®^ [poly(ICLC)] as i.n. prophylaxis prior to challenge with mouse-adapted SARS-CoV in Balb/c mice, where treatment had to be initiated within 8 h after virus challenge to prevent disease and death (31). The requirement for pretreatment with CAF09b indicates that the antiviral environment in the nasal tissue and lungs induced by i.n. administration of CAF09b, must be present at the time of infection. Possibly, virus-induced inhibition of IFN-I responses may hamper the effect of CAF09b, when administered post challenge during e.g. influenza- and coronavirus infections, where the innate response is corrupted by the virus (26, 32), whereby initial virus growth is allowed without immune pressure. This leads to delayed immune reaction to the infection and more severe disease.

The presented approach, may offer a means to tackle the ever-present threat of emerging viruses with pandemic potential, by offering a strategy to delay virus spread until vaccine roll-out can be initiated. Future studies will aim at testing longevity of protection against different viruses.

## CONCLUSION

These encouraging early preclinical data using the vaccine adjuvant CAF09b suggest a great potential for a pan-viral prophylaxis strategy against viral infections involving activation of innate immunity, especially IFN-I responses to establish anti-viral innate immunity against pandemic viruses. It will furthermore have the potential to give the immune system time to form the necessary adaptive immunity to protect against recurrent infections or even accelerate the development of adaptive immunity among infected individuals.

## ACKNOWLEDGEMENTS

We wish to acknowledge the excellent technical assistance received from Rune Fledelius Jensen and Julia Sid Hansen at the Department of Infectious Disease Immunology at Statens Serum Institut, and the Department for Biological Services at Statens Serum Institut.

## AUTHOR CONTRIBUTIONS

DC, GKP and FF conceived the idea for the concept and defined the project. DC, GKP, STS and JZ designed the experiments. GKP, STS, JZ, RJC, FZ and RT helped conducting the experiments. DC, GKP, STS and JZ wrote the manuscript.

