## Supplementary figures and images for "A novel Pan-viral prophylaxis strategy using vaccine adjuvant CAF09b protects against influenza virus infection"

### Supplementary figure 1

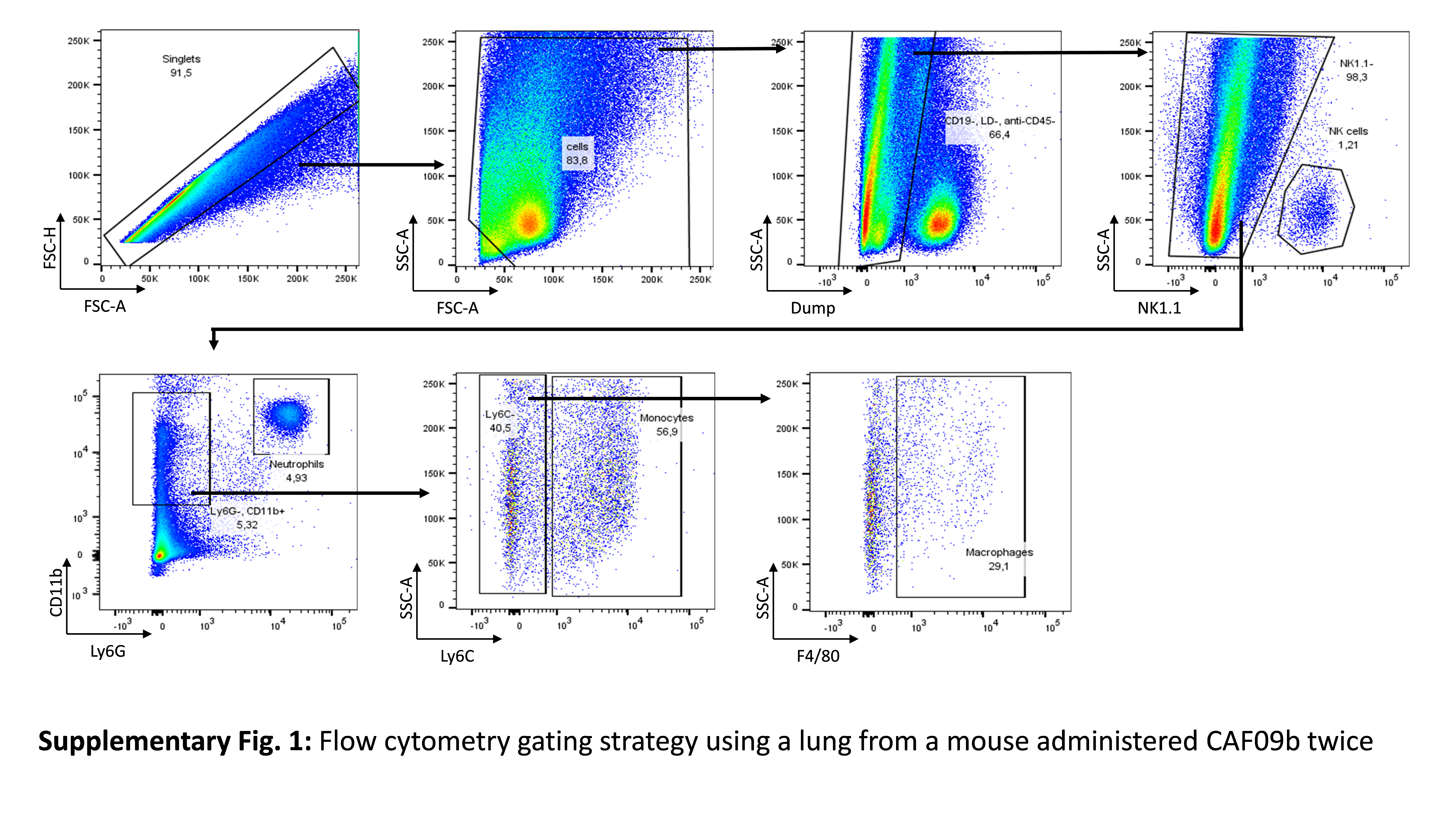

### Supplementary figure 2

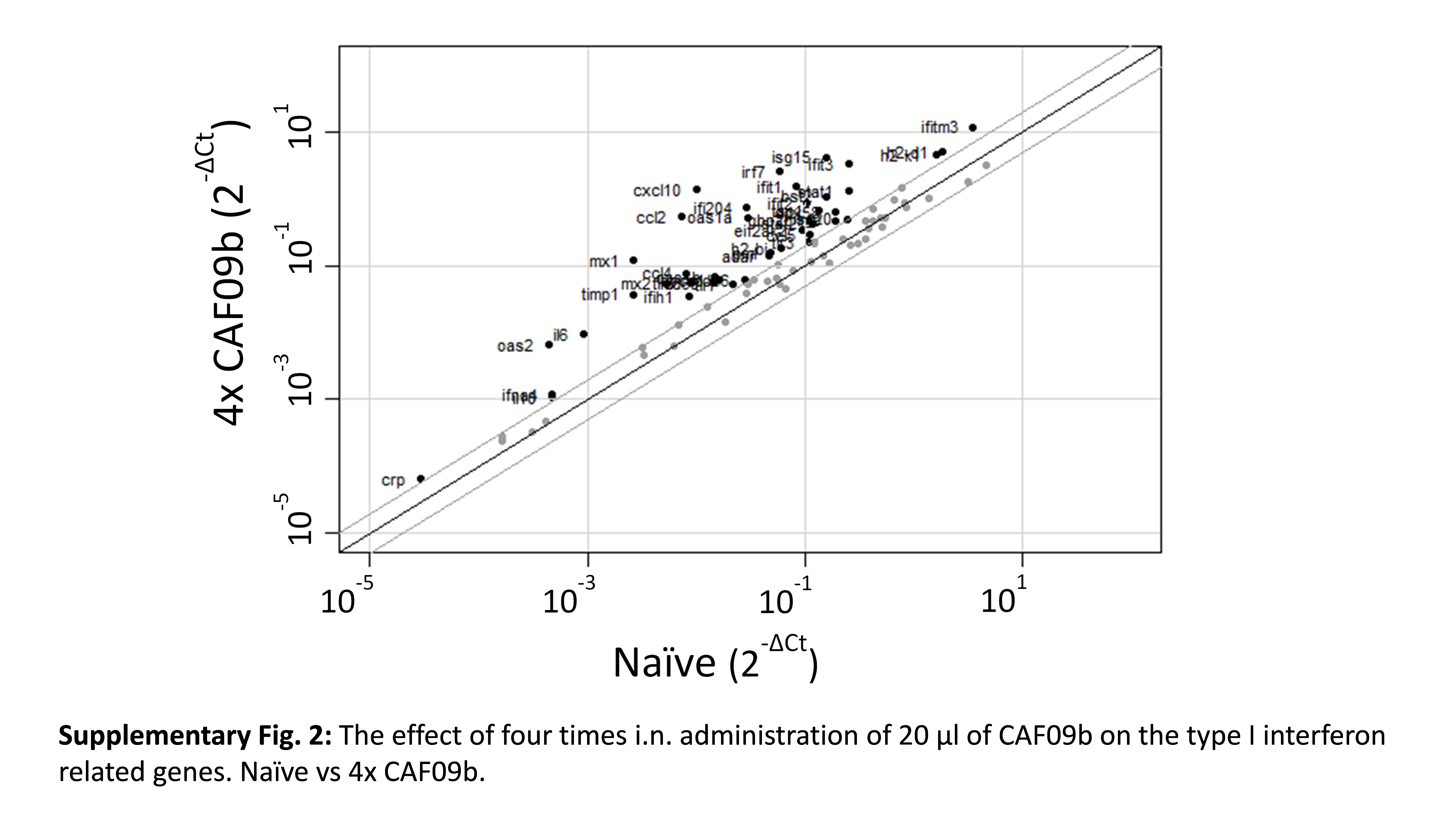

### Supplementary figure 3

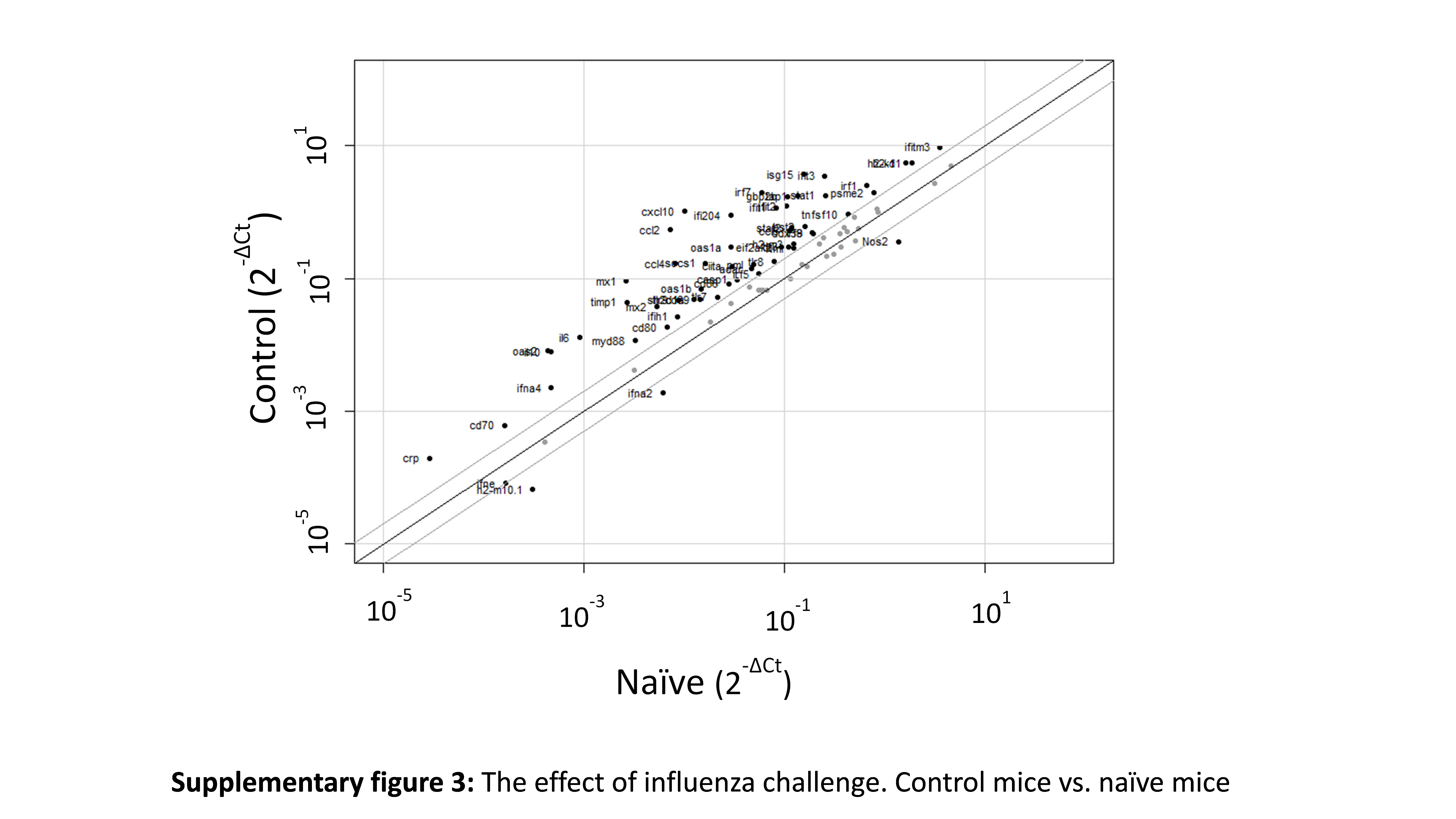
